# Blunted anterior midcingulate response to reward in opioid users is normalized by prefrontal transcranial magnetic stimulation

**DOI:** 10.1101/2024.10.03.616476

**Authors:** Kathryn Biernacki, Rita Z. Goldstein, Malte R. Güth, Nelly Alia-Klein, Suchismita Ray, Travis E. Baker

## Abstract

**Introduction:** Abnormalities in goal-directed behavior, mediated by mesocorticolimbic reward function and structure, contribute to worse clinical outcomes including higher risk of treatment dropout and drug relapse in opioid users (OU).

**Material and Method:** In a sham-controlled randomized study design, we measured whether robot-assisted 10Hz transcranial magnetic stimulation (TMS) applied to the prefrontal cortex was able to modulate anterior midcingulate cortex (MCC) electrophysiological response to rewards, in OU and matched healthy controls.

**Results:** We show that OU exhibit a blunted anterior MCC reward response, compared to healthy controls (t(39) = 2.62, p = 0.01, d = 0.84), and that this is normalized following 10-Hz excitatory TMS (t_(36)_ = .82, p = 0.42, d = 0.17).

**Conclusions:** Excitatory TMS modulated the putative reward function of the MCC in OU. Further work with increased sample sizes and TMS sessions is required to determine whether restoring MCC reward function increases reward-directed behaviors, which may enhance treatment success through the maintenance of treatment goals.

---

Opioid use disorder represents the pathological usurpation of the mesocorticolimbic reward system that normally serves several goal-directed processes (e.g., action selection and monitoring, reward valuation, decision-making)[1]. In support, a comprehensive meta-analysis indicated that opioid users (OU) demonstrate severe decision-making deficits, which have been linked to higher treatment dropout and rates of relapse, persisting years after cessation of use[2]. However, treatments for current or past opioid addiction fail to address such neurocognitive deficits directly, likely due to the limited extant circuit–based treatments (e.g., with transcranial magnetic stimulation, TMS) and neurocognitive measures (e.g., with EEG, fMRI) that are yet to be fully integrated as outcome measures into clinical trials[3]. Leveraging the significant translational potential of non-invasive brain stimulation[4], we report a combined EEG and TMS study, in a randomized sham-controlled design, aimed to quantify and modulate the reward function of the anterior midcingulate cortex (MCC) in OU.

The mesocorticolimbic circuit encompasses dopaminergic neurons that project from the ventral tegmental area (VTA) to various subcortical and cortical areas, including the MCC, where the VTA provides the main dopaminergic input[1, 5]. These dopamine neurons encode a reward prediction error (RPE) – the difference between predicted and received rewards – and current thinking holds that the MCC utilizes these RPEs to learn the value of rewards for the purpose of selecting the most appropriate action plan directed towards goals[6–8]. In this way, valuation by MCC appears to increase the motivation and effort to work for the reward (‘wanting’), as distinct from the hedonic enjoyment (‘liking’) of the reward when consumed[6, 9]. In humans, the impact of RPE signals on the MCC can be reliably measured using the reward positivity, an event-related brain potential (ERP) that is sensitive to rewarding feedback received during decision-making tasks[10, 11]. Converging evidence from multiple sources indicate that the reward positivity is produced in the MCC[6, 7], whereby simultaneous recording of EEG/fMRI signals show both a robust reward positivity and enhanced reward-related BOLD response in the MCC and striatum (Figure 1A), and computational work has confirmed that the reward positivity amplitude behaves as an RPE signal[12, 13]. Together, this evidence underscores the sensitivity of the reward positivity to mesocorticolimbic reward circuit activation.

**Figure 1.**
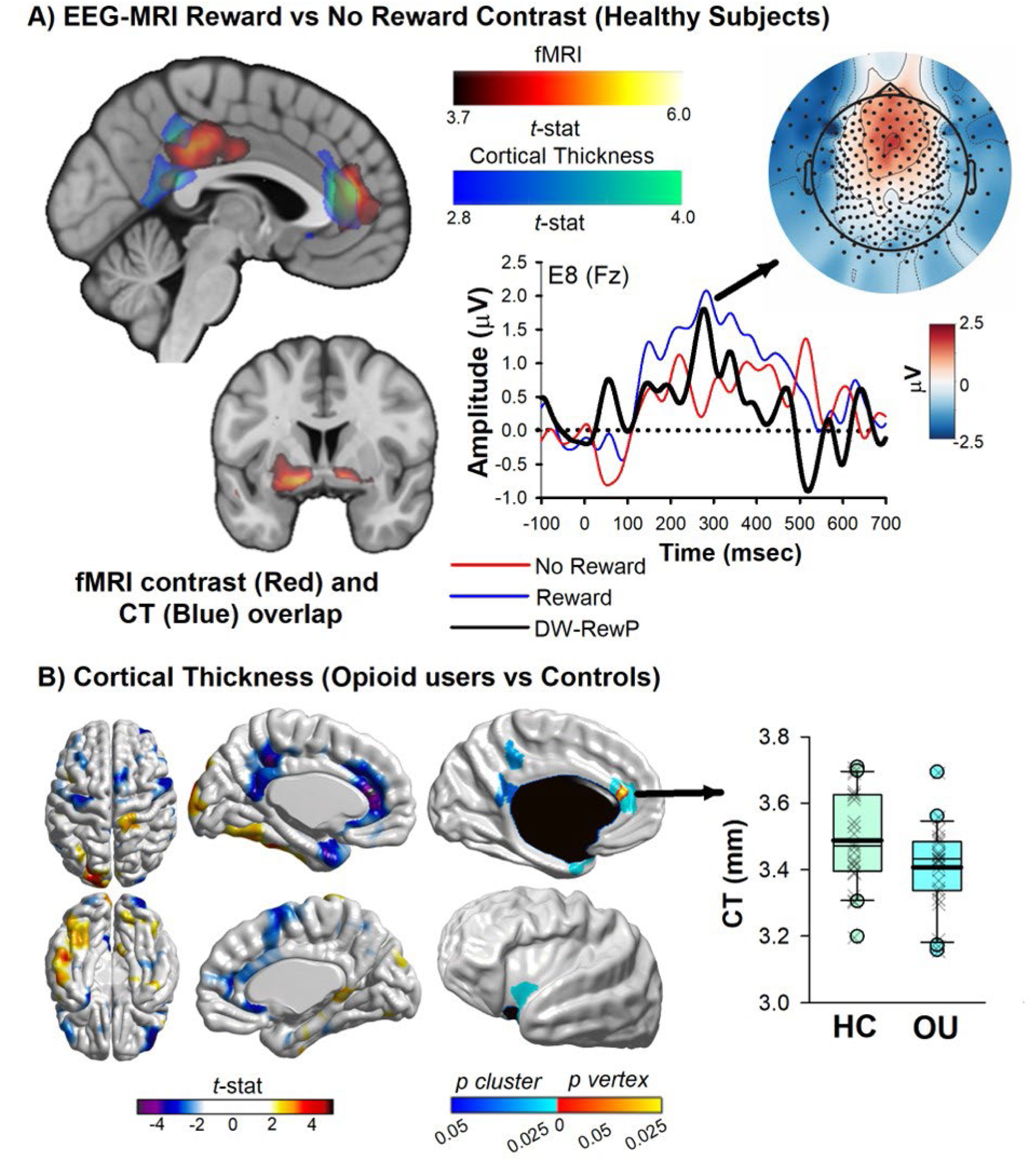
**A)** Simultaneous EEG-fMRI data in healthy controls (re-analysis of data from Güth et al., 2024). Left plot shows whole brain contrast between reward vs. no reward feedback in the virtual T-maze task. Clusters of increased reward-related BOLD signals identified in bilateral striatum, bilateral anterior midcingulate cortex, left posterior cingulate, and left dorsolateral prefrontal cortex (p < 0.05, FWE-corrected). Spatial overlap between cortical thinning in opioid users (OU) (Blue heat map) and fMRI reward clusters (Red heat map). Right plots show ERPs elicited by reward (blue), no reward (red) feedback and associated difference wave (black-reward positivity) at channel Fz. Scalp voltage maps associated with peak latency of the reward positivity. The reward positivity elicited in the T-maze task was clearly evident in the difference wave (mean = 3.00 μV, SEM = 0.52) peaking 260 ms after feedback onset, and was significantly different from zero, t(24) = 5.7, p < 0.001, 95% confidence interval = [1.9 μV, 4.07 μV]. **B)** MRI-based cortical thickness estimates (re-analysis of data from [21]. Left panel: t-statistic comparing OU to healthy controls (HC) at each cortical vertex. Right Panel: thresholded map showing areas of significant differences and cluster-corrected p-value in blue (p < 0.05, FDR-corrected). Clusters of cortical thinning in OU were identified in parts of the left anterior midcingulate cortex, left insula, left precuneus region, and left retrosplenial / posterior cingulate cortex (full details in SOM). The box plot denotes cortical thickness at the peak voxel within the MCC cluster between healthy controls and OU. The boundary of the box closest to zero indicates the 25th percentile, a thin line within the box marks the median, the thick line within the box marks the mean, and the boundary of the box farthest from zero indicates the 75th percentile. Whiskers (error bars) above and below the box indicate the 90th and 10th percentiles, and symbols denote outlying points outside the 10th and 90th percentiles. For full methods and results of existing datasets, please see Supplementary Material.

Chronic opioid use dysregulates and damages the VTA at the synaptic, cellular, and structural level[1, 14, 15], accompanied by dysregulated dopaminergic function[16] and altered functional connectivity among its various connected nodes[17]. Structural and functional abnormalities of the MCC, constituting one of these nodes, have indeed been reported in opioid use disorder, revealed using behavioral, electrophysiological, and neuroimaging tools[18–20]. For example, MRI-based cortical thickness estimates of a previously collected dataset[21] revealed that the MCC exhibited the greatest cortical atrophy in OU relative to healthy controls (Figure 1B). Dysregulated dopamine signaling in the MCC has also been observed in this population[22–25]. Moreover, individuals with substance use disorder produce a severely blunted reward positivity to monetary rewards[9, 26, 27], a finding which dovetails with previous neuroimaging studies implicating abnormal functioning of the cingulate cortex in addiction[28–31]. Together, this evidence suggests that MCC dysfunction would underlie several neurocognitive deficits (encompassing RPEs) and substance-related problems observed in OU, and therefore presents an opportunity for intervention, here a non-invasive brain stimulation with TMS.

While the MCC is too deep for TMS’ direct reach, neuroimaging studies have demonstrated that TMS can exert distant effects (on the MCC) if targeting interconnected regions[32]. Dense anatomical connections between the dorsolateral prefrontal cortex (DLPFC) and MCC[33–35] have therefore designated the DLPFC as a common TMS target, serving as an access node to the cingulate cortex. Notably, in positron emission topography studies, 10-Hz TMS over the left DLPFC, but not the right DLPFC, induced a significant increase in dopamine release in the cingulate cortex and striatum[36–38], attributed to regulation of activity of dopaminergic neurons in the VTA and its projection targets[36] and supported by previous animal work demonstrating that stimulating glutamatergic projections from the prefrontal cortex can regulate activation of dopaminergic neurons in the VTA[4, 36, 38]. Other mechanisms may be more direct and mediated by local activation of the neuron terminals in the DLPFC and MCC[33, 39]. For example, blood flow responses in the human MCC and evoked field-potential responses in the rodent cingulate are enhanced following TMS of the DLPFC, an effect proposed to reflect heightened neuronal excitability and lasting synaptic efficacy in the MCC[33]. Consistent evidence is provided by several fMRI studies demonstrating that TMS to the DLPFC induces a direct increase in brain hemodynamic responses in the cingulate cortex in both healthy and substance using populations [40–42]. We recently demonstrated that applying 10-Hz repetitive TMS to the left DLPFC increased the amplitude of the reward positivity in nicotine-deprived smokers[43] and polysubstance users[27], as well as improved decision-making capacity in polysubstance users[27] and healthy controls[44]. These findings bolstered our decision to use TMS of the DLPFC in treating reward-related MCC dysfunction in OU.

In this study, we examined whether robot-assisted 10-Hz repetitive TMS over the left DLPFC can modulate reward positivity amplitude in OU. To note, robot-assisted TMS has recently emerged as a superior alternative to conventional TMS methods because it allows real-time tracking of, and adjustment for, head movements, thereby providing increased reproducibility and accuracy of procedures[45, 46], especially in clinical populations where involuntary/excessive movements may be problematic[47]. First, we hypothesized that OU would demonstrate a blunted reward positivity amplitude, relative to healthy controls. Second, we predicted that Active 10-Hz TMS would normalize the reward positivity in OU randomized into the Active condition, such that they would produce a reward positivity comparable to that of healthy controls.

## Methods

### Participant recruitment and assessment

Participants were recruited through the Rutgers Alcohol and Drug Assistance Program, and local advertisements placed in the Newark community. Eligible participants were scheduled to complete one experimental EEG-TMS session. On the day of testing, participants were asked to provide informed consent and were then randomly assigned to either a TMS or Sham session. All subjects met the following inclusion criteria: English speakers; males and females aged 18-55 years old; ability to provide informed written or verbal consent. Exclusion criteria for all subjects included un-correctable visual impairment, uninterruptable central nervous system medication, severe brain injury (traumatic or acquired), and TMS contraindications (e.g., pregnancy, braces, history of seizures, medication which lowers the seizure threshold). Participants were financially reimbursed for their time and all data obtained was kept strictly confidential. The study was approved by the local research ethics committee and was conducted in accordance with the ethical standards prescribed in the 1964 Declaration of Helsinki.

In line with our previous work[27], problematic substance use was measured by the Alcohol, Smoking and Substance Involvement Screening Test (ASSIST)[48, 49]. To note, the ASSIST provides both a Global Continuum of Substance Risk (GCR) score^1^ and substance-specific sub-scores that identify the degree of problematic substance use for individual substances^2^. People were included in the chronic opioid-using group (OU) if they reported current or past problematic opioid use (e.g., heroin, oxycodone, hydrocodone, fentanyl) for at least 1 year, at least weekly use and/or met current criteria for opioid dependence according to the opioid-specific sub-score of the ASSIST (ASSIST score >12). People who reported past problematic opioid use had been abstinent from illicit opioid use for at least two weeks (average abstinence = 31.25 months, range = 0.5 to 240 months]). People were included in the healthy control group (HC) if they had no significant history of opioid use (ASSIST score <=3), or other drug-types (ASSIST score < 11 for each drug category or GCR <16). Additionally, controls and opioid users were excluded for severe mental health issues (e.g., schizophrenia, bipolar). Given the comorbid nature of drug addiction, we collected assessments of depression (Becks Depression Inventory) and anxiety (Becks Anxiety Inventory) to test for within group differences (Active vs. Sham, and OU vs. HC) (see Table 1).

**Table 1.** Sample characteristics.

| **Demographics** |  |  |  |  |  |
| --- | --- | --- | --- | --- | --- |
|  | **Opioid Use Disorder** |  | **Healthy Controls** |  | **OUD vs HC** |
|  | *Active* | *Sham* | *Active* | *Sham* |  |
| *N* | 16 | 18 | 22 | 24 |  |
| Age (years) | 42 (10) | 46 (10) | 37 (12) | 34 (11) | $p < 0.001$ |
| Sex (male/female/other) | 11/5/0 | 12/6/0 | 10/12/0 | 12/11/1 |  |
| Race (*n*) |  |  |  |  | n.s. |
| Black/African American | 10 | 12 | 7 | 11 | n.s. |
| White/Caucasian | 4 | 6 | 10 | 8 |  |
| Asian | 1 | 0 | 2 | 2 |  |
| Other or More Than One Race | 1 | 0 | 3 | 3 |  |
| Education (years) | 12 (2) | 13 (3) | 15 (2) | 15 (3) | $p < 0.001$ |
| rMT<sup>a</sup> | 72 (7) | 76 (4) | 72 (8) | 71 (10) | n.s. |
| TMS side effects | 1.1 (.03) | 1.0 (.06) | 1.1 (.02) | 1.1 (.02) | n.s. |
| **Opioid Use Behavior** |  |  |  |  |  |
| Opioid craving status (*n*)<sup>b</sup> |  |  |  |  |  |
| Pre-TMS | 3.3 | 3.0 | 1.1 | 1.1 | $p < 0.001$ |
| Post-TMS | 3.1 | 2.9 | 1.1 | 1.1 | $p < 0.001$ |
|  | n.s. | n.s. |  |  |  |
| Frequency opioid use (*n*)<sup>c</sup> |  |  |  |  |  |
| Weekly | 1 | 4 |  |  |  |
| Daily | 15 | 14 |  |  |  |
| Years opioid use | 13 (10) | 18 (11) |  |  |  |
| Opioid-dependence risk score<sup>d</sup> | 18 (14) | 21 (13) | 0 (0) | 0 (0) | $p < 0.001$ |
| **Other Drug Use Behavior** |  |  |  |  |  |
| Other drug risk scores<sup>d</sup> |  |  |  |  |  |
| Alcohol | 6 (6) | 9 (9) | 4 (6) | 5 (5) | $p = 0.012$ |
| Cannabis | 8 (9) | 6 (6) | 1 (3) | 1 (3) | $p < 0.001$ |
| Cocaine | 4 (8) | 8 (9) | 0 (1) | 0 (0) | $p < 0.001$ |
| GCR | 53 (26) | 73 (35) | 13 (19) | 10 (13) | $p < 0.001$ |
| Smoking (FTND) | 2 (2) | 4 (2) | 1 (2) | 1 (3) | $p < 0.001$ |
| **Mental Health** |  |  |  |  |  |
| Depression (BDI-II)<sup>e</sup> | 14 (11) | 16 (14) | 8 (8) | 10 (12) | $p = 0.014$ |
| Anxiety (BAI)<sup>f</sup> | 9 (11) | 7 (8) | 7 (8) | 13 (12) | n.s. |

<sup>a</sup> TMS-related side effects were evaluated by a review of symptoms (ROS) questionnaire at the end of the session.
<sup>b</sup> Opioid Craving Questionnaire
<sup>c</sup> Includes both illicit (e.g., heroin) and licit but abused opioids (e.g., oxycodone, fentanyl, hydrocodone).
<sup>d</sup> ASSIST substance specific scores; scores of 0-3 indicate low risk levels for substance dependence, 4-26 moderate risk, 27+ high risk; for alcohol, 0-10 indicates low risk, 11-26 moderate risk, and 27+ high risk.
<sup>e</sup> Beck Depression Inventory (BDI-II). Depression severity cut-offs for the BDI-II are as follows: 0–13 minimal, 14–19 mild, 20–28 moderate, and 29–63 severe.
<sup>f</sup> Beck Anxiety Index – Trait (BAI). Anxiety severity cut-offs for the BAI are as follows: 0–7 minimal, 8–15 mild, 16–25 moderate, and 26–63 severe.
\*To note there were no significant differences between variables within the OUD group

The final sample consisted of 34 participants in the OU group and 46 participants in the HC group (see Table 1 for full participant characteristics). Subjects were randomized into either Active (OU = 16; HC= 22) or Sham TMS groups (OU = 18; HC = 24). Two subjects were excluded due to recording issues (see EEG methods below).

### Electrophysiological Task: Virtual T-Maze

The reward positivity was measured using the virtual T-maze (see Figure 2), a reward-based choice task that elicits robust reward positivities[27, 50]. Participants navigate the virtual T-maze by pressing left and right buttons corresponding to images of a left and right alley presented on a computer screen. After each response, an image of the chosen alley appears, followed by a feedback stimulus (apple or orange) indicating whether the participant received 0 or 5 cents on that trial; unbeknown to the participants, the feedback is random and equiprobable. The experiment consisted of four blocks of 100 trials each separated by rest periods.

**Figure. 2.**
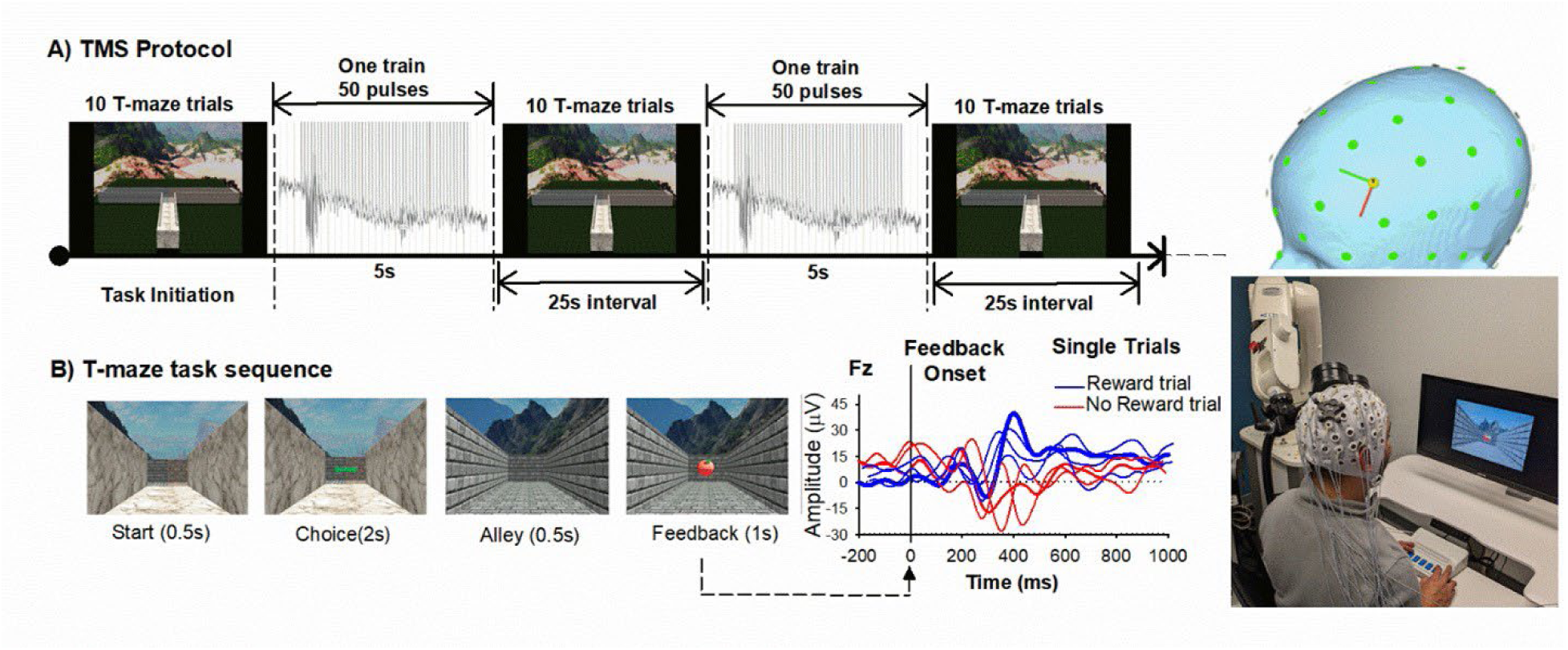
TMS Protocol displaying **A) TMS | T-maze** Block sequence and **(B)** single trial sequence. Right panels illustrate trial-to-trial ERPs associated with Reward (blue) and No reward (red) feedback, the robot-guided TMS set-up, 3D head view depicting the TMS target (electrode position F3).

### EEG Data Acquisition

EEG was recorded using a montage of 23 electrodes placed according to the extended international 10–20 system[51]. Signals were acquired using Ag–AgCl ring electrodes mounted in a nylon electrode cap with a conductive gel (Falk Minow Services, Herrsching, Germany). Signals were amplified by low-noise electrode differential amplifiers with a frequency response of DC 0.017–67.5 Hz (90-dB octave roll-off) and digitized at a rate of 1000 samples per second. Digitized signals were recorded to disk using Brain Vision Recorder software (Brain Products GmbH, Munich, Germany). Electrode impedances were maintained below 10 kΩ. Two electrodes were placed on the left and right mastoids, and EEG was recorded using the average reference. For the purpose of artifact correction, the horizontal EOG was recorded from the external canthi of both eyes, and vertical EOG was recorded from above the middle of the right eyebrow and electrode channel Fp2.

### Robot-guided transcranial magnetic stimulation

A MagPro X100 with the Cool-B70 figure-of-eight coil (MagVenture, Falun, Denmark) was used as the stimulator device. Following EEG-cap set-up, resting Motor Threshold (rMT) measurements were measured via visual twitch on the contralateral (right) hand. In particular, the coil was positioned over the supposed left motor cortex area (electrode location C3) and the coil was moved in a grid like search pattern until the location at which a reproducible abductor pollicis brevis response was detected. rMT was defined as the lowest stimulation intensity, expressed as a percentage of max output of the Magstim equipment, that reliably yielded a visible muscle twitch in the hand when stimulating the hand area of the contralateral motor cortex with a single pulse (3-5 times). Stimulation intensity for each experiment was set at 110% of maximal stimulator output. Participants in the sham condition were also measured for rMT (for blinding purposes), but did not receive active stimulation for the T-Maze task. Following rMT determination, a 3D model of the subjects’ head surface was created, and the robotic arm, head model, and the tracking system was registered to a common coordinate system (SmartMove, ANT Neuro, Enschede, The Netherlands). The TMS target was then placed on the x y z coordinate corresponding to electrode location F3, and the corresponding point on the surface was selected and the orientation of the coil was calculated tangentially to cortical surface and 45° to a sagittal plane based on the head model. The virtual coil position was then transformed to robot coordinates, which defined the movement of the robot to the corresponding target position relative to the subject’s head. The position of the cranium was continuously tracked, and the trajectory of the coil’s path was adapted to head movements in real-time. This procedure guaranteed a precise <1 mm coil-to-target position and allowed free head movements during the experiment.

### Design

Using a randomized control between-subjects design, participants were randomized into either the TMS Active or Sham group (coil flipped 180° to mimic auditory stimulation) during task performance. Participants were blinded as to the condition during the experiment, and received a debriefing statement at the completion of the experiment detailing which group they were randomized to. All participants completed consent protocols, and questionnaires and were then fitted with the EEG cap for ERP recording. Following the electrode cap set-up and rMT measurements, the robotic arm positioned the TMS coil over the electrode location F3, and maintained the coil position 10 mm from the scalp at a 45°angle (Figure 2). The EEG was recorded continuously while participants freely navigated the virtual T-maze to find monetary rewards (see Figure 2B) and participants were asked to respond in a way that maximized their rewards. The experiment consisted of four segments of 100 trials, each consisting of 10 blocks of 10 trials separated by rest periods (i.e., 4 segments × 10 blocks × 10 trials for 400 trials total; 200 reward and 200 no reward trials; Figure 1C). From the start of the T-Maze task, participants received 50 rTMS pulses delivered at 110% at 10-Hz continuously over the predefined left DLFPC target (duration: 5 seconds) immediately before each block of 10 trials (duration: 30 seconds per block). The duration between the last pulse of the train and first trial of each block exceeded 100 msec, allowing EEG data to be collected without TMS artifacts. The total duration of the task lasted approximately 20 minutes (2000 TMS pulses). The Sham condition received the same protocol, however, the coil was flipped 180° to ensure that participants did not receive active simulation but received the same auditory sensation. No subjects reported any side effects during or after the real or sham TMS (headache, nausea, etc.).

### Data Processing and Analysis

EEG post-processing and data visualization were performed using Brain Vision Analyzer software (Brain Products GmbH). The digitized signals were filtered using a fourth-order digital Butterworth filter with a bandpass of .10–20 Hz. A 1000-msec epoch of data extending from 200 msec before to 800 msec after the onset of each feedback stimulus was extracted from the continuous data file for analysis. Ocular artifacts were corrected using the eye movement correction algorithm described by Gratton, Coles, and Donchin [52]. The EEG data was re-referenced to linked mastoids electrodes, and data was baseline-corrected by subtracting from each sample the mean voltage associated with that electrode during the 200-msec interval preceding stimulus onset. Muscular and other artifacts was removed using a ±150-μV level threshold and a ±35-μV step threshold as rejection criteria. ERPs were then created for each electrode and participant by averaging the single-trial EEG according to feedback type (Reward, No reward).

The reward positivity was identified following a standard difference wave approach by subtracting the average reward waveform from the average no reward waveform for every electrode and participant, resulting in an average difference wave (i.e., the reward positivity). This “difference wave” approach, which was recommended in a meta-analysis of reward positivity studies [12], is a common analysis method used in ERP research to isolate neural processes by removing sources of variance that are common to both conditions. Because positive RPEs (indicating that ongoing events are better than expected) are associated with large reward positivities, and negative RPEs (indicating that ongoing events are worse than expected) are associated with the absence of the reward positivity, the difference wave approach eliminates neural activity that is common to both positive and negative feedback stimuli under the assumption that the underlying neurocognitive processes interact linearly. In other words, the difference wave isolates the reward positivity from other ERP components and provides an indirect measure of the positive RPE signal embedded in the ERP waveform. Here, the reward positivity amplitude was determined by identifying the maximum absolute amplitude of the difference wave within a 200- to 400-ms window following feedback onset, and evaluated at front-central electrode Fz where it reached its maximal.

## Results

### Reward Positivity

A two-way ANOVA assessed the interaction between TMS condition (Active vs. Sham) and Group (OU vs. HC) on reward positivity amplitude (covariates included age, sex, and global drug use score). In line with our hypothesis, this analysis revealed a main effect of TMS, F_1,72_ = 5.5, p = .02, η^2^ = .07, quantified by an interaction between TMS and Group, F_1,72_ = 6.9, p = .01, η^2^ = .09 (Figure 3 A-C). The Group main effect did not reach significance, F_1,72_ = 1.4, p = .24, η^2^ = .01. Follow-up analyses of the TMS main effect showed that the Active TMS group displayed a larger reward positivity (mean = 4.85 μV, SEM = 0.44) compared to the Sham TMS group (mean = 3.38 μV, SEM = 0.44), p = .02, d = 0.54. Follow-up analyses of the interaction showed two key findings. First, within the Sham TMS group, OU exhibited a blunted reward positivity to monetary feedback (mean = 1.98 μV, SEM = 0.72) when compared to HC (mean = 4.77 μV, SEM = 0.65), t_(39)_ = 2.62, p = 0.01, d = 0.84. Second, the delivery of Active 10-Hz TMS normalized this impairment. Specifically, no differences in reward positivity amplitude were observed between the Active OU group (OU: mean = 5.13 μV, SEM = 0.72) and HC groups (Active HC: mean = 4.56 μV, SEM = 0.64), t_(36)_ = .82, p = 0.42, d = 0.17 | Sham HC (t_(37)_ = −.72, p = 0.48, d = 0.11; see Figure 3D). The Active OU group also displayed a larger reward positivity compared to the Sham OU group, t_(32)_ = −3.2, p = 0. 003, d = .98. No differences were observed between Active and Sham HC (p > .05) groups.

**Figure 3.**
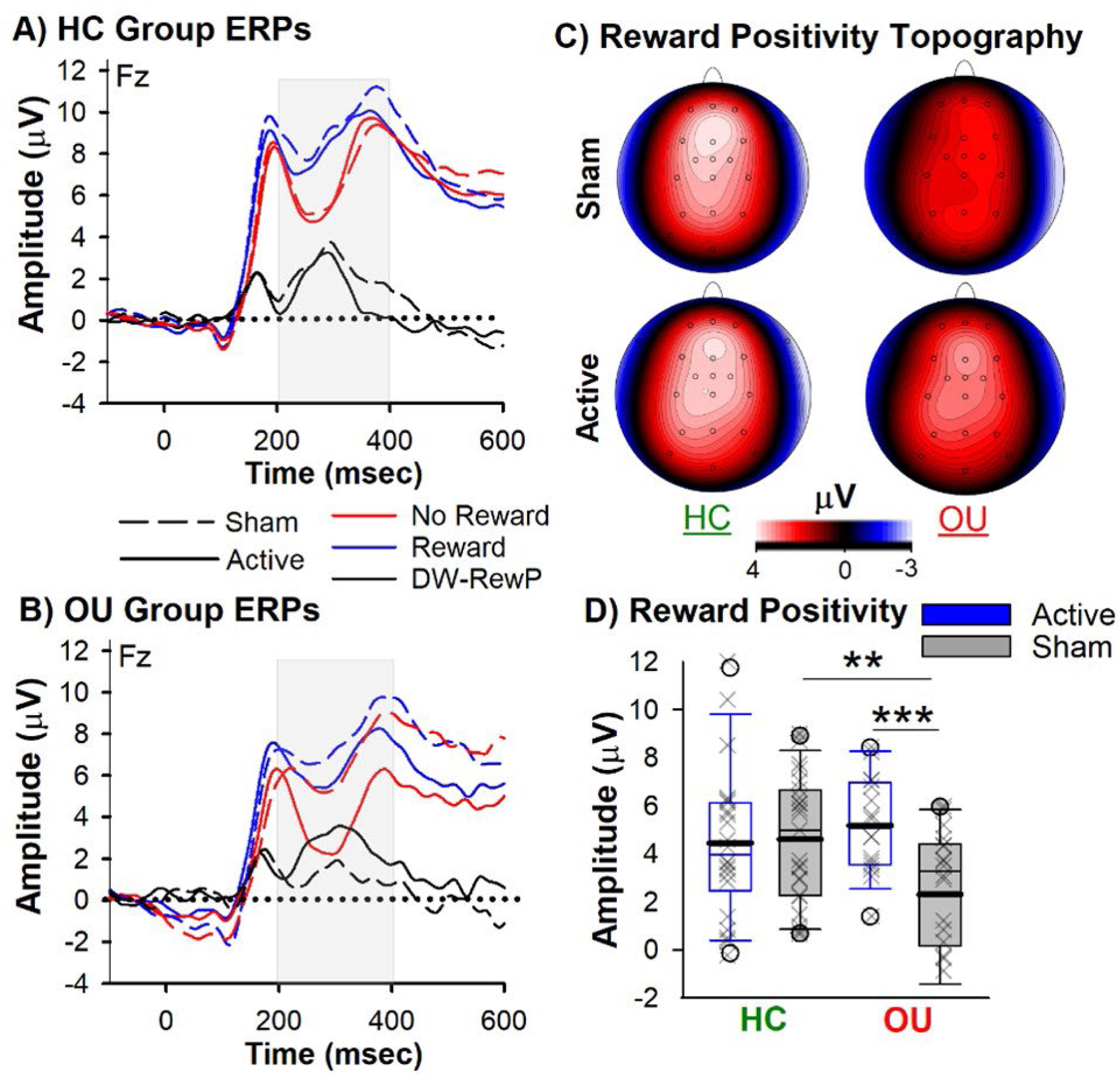
Reward Positivity Results. Grand-Averaged ERPs elicited by reward (blue) and no reward (red) feedback, and associated difference wave (black-reward positivity) for **(A)** Healthy Controls [HC] and **(B)** Opioid Users [OU] groups (Solid lines: Active TMS, Dashed lines: Sham). **(C)** Topoplots illustrate the reward positivity peak. **(D)** The box plot denotes reward positivity amplitude for the Sham (grey bars) and Active (blue bars) HC and OU groups. The boundary of the box closest to zero indicates the 25th percentile, a thin line within the box marks the median, the thick line within the box marks the mean, and the boundary of the box farthest from zero indicates the 75th percentile. Whiskers (error bars) above and below the box indicate the 90th and 10th percentiles, and symbols denotes outlying points outside the 10thand 90th percentiles. Significant effects are shown as follows: **p < .01, ***p < .005 (two-tailed).

### P200 and P300

The amplitudes of other prominent ERP components (P200 and P300) were equivalent between the TMS conditions and Groups (p > .05), confirming that the TMS effect of interest and group differences were isolated to the predicted ERP component, the reward positivity, and did not reflect an overall processing or attentional difference to task demands across sessions. Interestingly, the amplitude of the N200 component to no reward feedback was reduced in the Sham OU group (mean = −3.57 μV, SEM = .62) compared to Sham HC (mean = −5.88 μV, SEM = .73: t(39) = 2.3, p = 0.02) and Active OU (mean = −6.04 μV, SEM = 1.0, t(32) = 2.1, p = 0.04) group. Further, within the Sham OU group the amplitude of the N2 to reward feedback mirrored the N200 to no reward feedback (p > .05), confirming that reward feedback failed to induce the reward positivity in these individuals, replicating previous results [9, 53]. In regards to the craving measures, no differences were observed between pre- and post-TMS opioid craving scores or within and between the Active and Sham OU groups (see Table 1), and no inter-relationships were observed with our ERP measures.

## Discussion

The MCC is one of the main cortical targets of the mesocorticolimbic reward system originating in the VTA, a critical neural locus for opioid reward and addiction[1, 16]. It has been argued that the enhancement of dopamine release triggered by opioid use can resemble a positive RPE: whereas the dopamine response to the reward itself disappears when the reward is predicted, transient increments in dopamine levels caused by the drug itself persist over time because they are pharmacologically mediated through direct action on dopamine neurons, and thus produce positive prediction errors even when drug-related events are expected[54]. By inducing continuing strong dopaminergic stimulation of VTA projected targets, such as the MCC[5], the augmented dopaminergic RPEs exaggerate the value of states that precede drug consumption and, in turn, desensitize the system to nondrug rewards[55]. In other words, by “hijacking” brain reward systems, drug-related behavior and cues become overvalued. Consistent with these arguments and theoretical models, our current electrophysiological data show that individuals with a history of opioid misuse exhibit a blunted (nondrug) reward positivity — an electrophysiological signal that indexes sensitivity of the MCC to positive RPE signaling — indicating that reward feedback failed to induce an MCC reward response in these individuals. We argue that such abnormal MCC reward functioning would underlie decreased valuation and effortful pursuit of a wide range of goal-directed behaviors and a marked narrowing of life goals to obtaining and using opioids[53].

To compound this impairment, we also show that the N200 component, a well-established index of cognitive control[56], was attenuated in Sham OUs relative to Sham controls. The N200 is a frontocentral negative deflection in the ERP elicited by unexpected, task-relevant events, including both unexpected positive and negative feedback[10, 57]. Empirical and theoretical work suggests that the neural generator of the N200 is the MCC[56, 58], the N200 amplitude reflects the level of effortful control applied by MCC over task performance[6], and the N200 provides a sensitive index of cognitive control deficits (e.g., inhibitory control) in substance use disorders [59, 60]. In line with this evidence, our findings of an attenuated N200 to no reward feedback in OU indicate that task-relevant events failed to elicit an MCC-dependent control process in these individuals. That the reward positivity and N200 can serve as independent measures of MCC processes of reward valuation and effortful control, respectively^3^, suggests that OU expresses impairments in both these functions, which is in line with the Impaired Response Inhibition and Salience Attribution (iRISA) model of addiction[29, 61, 62]. The iRISA model proposes that dysregulated networks involved in salience attribution (the reward network) and inhibitory control (the executive network) underlie drug seeking and taking, such that addicted individuals increase recruitment of these cognitive processes during the pursuit of drug-related goals, but a blunted response during non drug-related goals, as we have shown here by an attenuated reward positivity and N200.

To counteract the aberrant MCC reward function in OU, we applied a robot-assisted TMS intervention used in our previous studies[27, 43]. Consistent with our previous findings, applying 2000 pulses of 10-Hz TMS to the left DLPFC normalized the MCC reward response, as revealed by a quantifiable increase in reward positivity amplitude in the OU group receiving Active vs. Sham TMS, and at par with the reward positivity in both the Active and Sham healthy control groups. Furthermore, we also observed an increase in N200 amplitude in the OU group receiving Active TMS compared to Sham OU group. Although the N200 result was not predicted, it does align with our previous study showing that 10-Hz TMS to the left DLPFC equivalently enhanced the amplitude of the N200 to reward and no reward feedback in nicotine users[43]. Current evidence suggests that stimulating the left DLPFC using high-frequency TMS (e.g., 10-Hz) can enhance dopamine release, neuronal activity, and cerebral blow flow in the cingulate cortex, a modulatory effect mediated by local coactivation of the DLPFC and MCC[33, 63], and/or indirect modulation of the VTA, resulting in an increase in dopamine transmission among its frontal–striatal targets[4, 36]. We propose that an TMS-induced increase in VTA dopaminergic activity heightened the positive RPE signal conveyed to the MCC during monetary reward trials, as revealed by a larger reward positivity in the Active OU group. Further, because ERPs are a spatiotemporally smoothed version of the local field potential integrated over the cortex[64], our observation of an TMS-induced increase in N200 amplitude to no reward feedback confirms that direct stimulation of the DLPFC-MCC pathway enhanced MCC excitability, and possibly the MCC control response during task execution.

To summarize, while theories of MCC function span a plethora of functions[65], we believe our findings are consistent with MCC’s role in the valuation and selection of effortful behavior[7, 8]. Accordingly, the MCC utilizes RPE signals (reflecting reward valuation) to assess how much control to allocate over task performance (reflecting effort expenditure), and that these factors manifest in two electrophysiological signals generated within MCC: the reward positivity and N200, respectively[6]. Put another way, the MCC function should underlie the ability to select and motivate effortful behaviors according to the value of rewards received during task execution. We propose that individuals with a history of opioid misuse produced a low reward value (blunted reward positivity) to small monetary incentives used in this task, which in turn resulted in reduced effortful control (reduced N200) during task performance. By extension, an unstable valuation and control function of MCC ⸺ along with structural loss to the MCC (see cortical thickness results) ⸺ would present itself as signature characteristics of several substance-related problems observed in current and past OU. To alleviate this impairment, it can be argued that active TMS of the left DLPFC facilitated the normalization of the MCC role in assignment of a high positive value to the pursuit of small monetary incentives received during the task (enhanced reward positivity), and as a result, increased effortful control over task performance (enhanced N200). To translate this to a clinical setting, modulating MCC putative function long-term could potentially help recovering OU maintain continued motivation by assigning sufficient value to self-directed, goal-driven behaviors including those aligned with treatment goals.

Of course, MCC constitutes only one node within several larger networks involved in the drug addiction cycle (onset of substance abuse, transition to addiction, chronic relapse), which raises the question: Do abnormal indicators of MCC activity in OU reflect intrinsic impairments to the MCC itself, or abnormal input to MCC from other brain areas, or both? For example, reduced reward positivity and N200 amplitude in OU could indicate malfunction of MCC proper (where these ERP components are generated), reduced ventral striatal and VTA activation (which is strongly correlated with reward positivity amplitude), and/or an inability to synchronizing its activity within a larger neural network[6, 56]. Although we cannot confirm such possibilities, our whole-brain cortical thickness estimates in a separate cohort of inpatients with opioid use disorder[21] indicated that both factors may be involved. In particular, the MCC demonstrated the greatest cortical atrophy in an opioid use disorder population and such structural abnormalities in MCC clearly point to local dysfunction. By contrast, several key cortical targets of the VTA, namely the insula, precuneus, and posterior cingulate cortex, also exhibited high cortical atrophy, which aligns with network dysfunction in addiction proposed by iRISA[29, 61, 62], as well as the network-spread hypothesis implicated in neurodegenerative disorders (i.e., disease progression follows the neuronal connectome of the brain)[66]. In support of this hypothesis, our previous cortical thickness estimates in Parkinson’s Disease indicated that regions exhibiting the most atrophy are part of a connectivity network centered around the substantia nigra, the neural loci of Parkinson’s Disease[67]. Because MRI/PET studies have shown that grey matter density in mesocorticolimbic regions, particularly the MCC, appears strongly correlated with dopamine receptor availability, suggesting that the denser the grey matter the more D_2_-like receptors it can support[68]. In line with our functional results, we attribute these structural results to a dysregulation of VTA dopaminergic neurons by opioid misuse[5, 69] and potentially the networks in which the VTA and MCC serve[40, 62].

We have posited a central role for the MCC action-selection mechanism in substance use disorder, and our findings indicate that 10-Hz TMS to the left DLPFC modulated this mechanism in OU. Another outstanding question is whether the postulated therapeutic effects of TMS are based on direct or indirect (network) effects on MCC function[70]? On the one hand, the observation of dense anatomical connections between the DLPFC and MCC has led to the proposal that TMS effects on MCC activity might be more direct[33]. On the other hand, DLPFC stimulation and subsequent activation of the VTA can lead to an indirect modulation of dopaminergic activity in the MCC[36]. In fact, we suspect that both factors may be involved. We speculate that such TMS may have potentiated positive RPE signaling and enabled the MCC to respond dynamically to both positive and negative RPE signals, thereby driving a larger differential response between good (positive RPE) and bad (negative RPE) outcome ERPs. Although we cannot confirm such possibilities, future investigations of the impact of TMS on electrophysiological activity of the MCC in substance use populations are warranted. For instance, if both TMS-induced mechanisms of action (i.e., dopaminergic and neuronal modulation) are involved in modulating MCC-related electrophysiological signals, then electrophysiological recordings in animal models of addiction will yield a comparable difference in RPE signals and evoked field-potential responses in the MCC following TMS.

Future research may address some of the study’s limitations. First, our subjects do not represent a sample of individuals with an opioid use disorder seeking treatment, and results remain to be replicated in these and other groups (e.g., long-term abstaining) of OU. Given the data demonstrating persistence with protracted abstinence from opioid use of both abnormal connectivity within the reward networks and decision-making impairments[2, 71–73], we speculate our results to replicate, with a meaningful clinical impact, across OU populations^4^. Second, given the stark difference in tactile sensation between sham and active TMS, future studies should consider adopting a double-blind procedure (e.g., using a placebo coil) to reduce participant and experimenter bias. In regard to clinical outcomes, previous studies have shown that one session or twenty daily sessions of 10-Hz TMS to the left DLPFC reduced cue-induced craving scores in heroin users[74, 75]. We failed to replicate the effect of TMS on pre- and post-craving scores, nor did we find an association between craving and our ERP data. Several factors could be at play, including diverse and small sample, sensitivity of the craving assessment, non-cue-reactivity task, and the number of TMS pulses and sessions. On the other hand, perhaps the impact of TMS on craving is small and unreliable[76], and future studies should focus more on neurocognitive outcomes[3] or targeting brain regions implicated in cue-induced craving processes, such as the hyperdirect pathway[77, 78] or insula[79]. Finally, while a good starting point, the conventional targeting approach used in the current study (scalp-landmark) frequently misses the DLPFC[32]. The DLPFC spans a large anatomical region and is highly variable across individuals[34, 80], thus simply placing the TMS coil on a DLPFC-based scalp landmark would be less than ideal for precision medicine. For example, we have previously shown that TMS responsiveness (as evaluated by the reward positivity) was associated with prefrontal cortical thickness, estimated connectivity tracts between the TMS target and MCC, as well as the striatum, and a greater left lateralization of the cingulum bundle[43]. These considerations highlight the need to optimize targeting techniques for DLPFC-MCC precision (e.g., individualized targets based on DLPFC structure, function, or connectivity with MCC) in order to increase the efficacy of TMS to normalize MCC activity long-term in substance use disorder [40].

## Conclusion

When applied in trains of repetitive pulses at around 5–10 Hz, TMS is thought to induce stable potentiation-type plasticity in the targeted circuit, potentially modifying activity in brain-wide networks [41, 81, 82]. Combined, our results suggest that, paired with conventional treatment programs and standard of care, modulating MCC putative function with TMS could potentially help recovering OU maintain continued motivation by assigning sufficient value to self-directed, goal-driven behaviors including those aligned with treatment goals. It remains to be tested whether modulating the MCC’s putative function in one session could be replicated across sessions and, importantly, extend beyond the laboratory to alleviate main addiction symptoms and improve clinical outcomes (e.g., enhance effortful pursuit of a wide range of goal-directed behaviors to improve treatment adherence, cognitive and social functioning, and employment rates). Nevertheless, by highlighting important opioid use disorder-relevant neurocognitive responses to TMS, this research may represent an important step in this promising direction.

## Supporting information

Supplemental methods and results

## Author contributions

T.E.B and K.B conceptualized and designed the research. K.B collected the data, and S.R. assisted with subject recruitment and assessment. K.B and T.E.B analyzed the EEG data, M.R.G and T.E.B collected and processed the EEG-fMRI dataset. N.A and R.Z.G collected the MRI dataset in the separate cohort of opioid use disorder patients and healthy controls, and T.E.B, and R.Z.G. analyzed the cortical thickness dataset. T.E.B and K.B wrote the manuscript, with suggestions by all authors.

## Acknowledgements

We thank Sally Cole, Galit Karpov and Mei-Heng Lin for their contribution to data collection, and Merna Zaki for aiding in cortical thickness data analysis.

## Conflict of interest

The authors report no conflicts of interests.

## Funding

This work is supported by the National Institute on Drug Abuse of the National Institutes of Health [Award Number 1R21DA049574-01A1] and the National Center for Complementary and Integrative Health [Award Number 1R01AT010627-01].

## Footnotes

1 According to cut-offs scores established in validation studies of the Alcohol, Smoking and Substance Involvement Screening Test (ASSIST), mid-range Global Continuum of Substance Risk (GCR) scores between 16 and 39 are an indication of hazardous or harmful substance use. GCR scores of 39.5 and higher suggests that the individual is at high risk of substance dependence (Humeniuk et al., 2008, Newcombe et al., 2005).

2 ASSIST substance-specific scores of 0-3 indicate low risk levels for substance dependence, scores 4-26 moderate risk, and scores 27+ as high risk; alcohol risk is defined at different levels, however, with scores 0-10 indicating low risk, 11-26 moderate risk, and 27+ high risk (Humeniuk et al., 2010).

3 Recent examinations of the reward positivity and the N200 have provided a nuanced account about their relationship, such that the N200 constitute a baseline control response by MCC to unexpected events and its amplitude is regulated up and down by positive and negative RPE signals conveyed to MCC. Thus, positive dopamine RPE signals conveyed to MCC about 300 msec after unexpected reward feedback drive a positive-going deflection in the ERP (the “reward positivity”) that cancels out or inhibits production of the N200. Because the variance in the amplitude of the N200 is driven more by unexpected rewarding events than by non-rewarding events, the reward positivity is typically operationalized as the size of the difference wave calculated between good (positive RPE) and bad (negative RPE) outcome ERPs (Holroyd et al., 2008; Baker and Holroyd, 2011; Hajihosseini and Holroyd, 2013).

4 Splitting the analysis between current and past opioid users does not change the main outcome: Active TMS group displayed a larger reward positivity (Current: mean = 5.90 μV, SEM = 0.71 | Past: mean = 4.73 μV, SEM = 0.67) compared to the Sham TMS group (Current: mean = 2.8 μV, SEM = 0.83, p<.05 | Past: mean = 1.66 μV, SEM = 1.2, p < 0.05).

## Notes

### Competing Interest Statement

The authors have declared no competing interest.

