## Supplemental methods and results for "Blunted anterior midcingulate response to reward in opioid users is normalized by prefrontal transcranial magnetic stimulation"

**Supplementary Material**

### Analysis of existing datasets

#### Cortical thickness analysis

We utilized an existing neuroimaging dataset to conduct a cortical thickness analysis between patients with opioid use disorder and healthy controls [1]**.** All participant demographic and clinical data are detailed in Ceceli et al., 2023 (Table 1). MRI scans were acquired using a Siemens 3.0 T Skyra (Siemens Healthcare) with a 32-channel head coil. T1-weighted anatomical image acquisition parameters were as follows: 3D MPRAGE sequence with 256 × 256 × 179 mm3 field of view, 0.8 mm isotropic resolution, repetition time/echo time/inversion time = 2400/2.07/1000 ms, 8° flip angle with binomial (1, −1) fat saturation, 240 Hz/pixel bandwidth, 7.6 ms echo spacing and in-plane acceleration (GRAPPA) factor of 2, with a total acquisition time of approximately 7 min.

Following our previous methods of cortical thickness analysis [2-5], all T1-weighted MRI images were processed using the CIVET pipeline (version 2.1) (www.bic.mni.mcgill.ca/ ServicesSoftware/ CIVET). Briefly, native T1-weighted MRI scans were corrected for non-uniformity artifacts using the N3 algorithm. The corrected volumes were masked and registered into stereotaxic space, and then segmented into gray matter, white matter, cerebral spinal fluid and background using an advanced neural net classifier. The white matter and gray matter surfaces were extracted and resampled to a stereotaxic surface template to provide vertex based measures of cortical thickness. For each participant, cortical thickness was then measured in native space using the linked distance between the two surfaces across 81924 vertices. Each subject's cortical thickness map was blurred using a 20-mm full width at half maximum surface-based diffusion smoothing kernel to impose a normal distribution on the corticometric data, and to increase the signal to noise ratio.

Statistical analyses were performed using SurfStat (www.math.mcgill.ca/keith/surfstat/), a statistical toolbox created for MATLAB (The MathWorks, Inc., Nathan, MA, USA). Each participant's absolute native-space cortical thickness was linearly regressed against group (OU and Healthy controls) and Time (Time 1 and Time 2: scans were separated by 3-4 months into inpatient treatment and equivalent times in HC) at each cortical point after accounting for the effects global brain volume, age, and sex. In summary, the following model was fitted to each one of the 81924 cortical points ($Y = b0 + b_i\,Group + b_{ii}Time + b_2GLOBAL + \mathcal{E}$) where Y = Cortical

Thickness, b0 = Y intercept, b2 = regression coefficients for effects of global thickness volume, ε = error term. For each cortical point, the coefficient of the Group and Time regressor was estimated and a resultant t-test value calculated, thereby producing a t-statistic map (Figure 1b). A t-statistic threshold of statistical significance was established, taking into account multiple comparisons via the false discovery rate (FDR) method. The resulting thresholded maps were projected on an average surface template for visualization (See Figure 1b).

**EEG-fMRI data acquisition**

To highlight reward-related activity in fMRI and ERP data, we utilized an existing neuroimaging dataset [6]. EEG data for all 256 channels were collected at a sampling rate of 1 kHz using an MR-compatible HydroCel Geodesic sensor net (Electrical Geodesics Inc., Eugene, USA). In addition to scalp electrodes, a four-lead electrokardiogram (EKG; two active, two dummy) was recorded for later artifact correction. One active EKG lead was placed on the lower end of the sternum and one below the left chest on the rib cage. All electrodes recorded from inside the scanner were online referenced to E257, which is the geodesic electrode placement system's equivalent to Cz in the international 10-20 system[7]. Data were recorded using the Net Station software (Version 5.4.2) applying an online bandpass filter excluding data above 100 Hz and below 0.001 Hz. All electrode impedances were kept below 50 kΩ. MRI data were collected in a 3T scanner (Trio Tim, Siemens) with a 12-channel head coil at the Rutgers Brain Imaging Center. Anatomical images were acquired with a T1-MPRAGE sequence, consisting of 176 sagittal slices (1 mm isotropic voxel; TR = 2500 ms, TE = 2.52 ms, flip angle = 9°). A dual-echo gradient-echo sequence was used to assess a B0 inhomogeneity gradient field map (TE1 = 5.19 ms, TE2 = 7.65 ms, TR = 400 ms). Functional whole-brain images were collected in an axial orientation by using a T2-weighted gradient echo planar imaging sequence (TR = 2000 ms, TE = 25 ms, flip angle = 90°) with an isotropic voxel size of 3.3 mm (64 x 64 matrix; 208 x 208 mm2 field of view). 35 slices per volume were acquired in an ascending, interleaved order. Functional image acquisition sessions lasted 15 minutes, resulting in 450 volumes for each participant.

**EEG data preprocessing and analysis**

EEG data concurrently collected with fMRI are affected by two major MRI-related artifacts: Gradient artifacts (GA) and ballistocardiac artifacts (BCA). Both average artifact subtraction for GA correction[8] and initial BCA correction based on principle component analysis[9] were carried out using Brain Vision Analyzer 2.1 (Brain Products GmbH, Gilching, Germany). Subsequent preprocessing including Independent Component Analysis (ICA), filtering, re-referencing, and segmentation were performed using custom scripts in Python 3.9 and MNE-python (Version 1.0.0) for M/EEG analysis[10]. Next, the EEG signals were filtered using a Butterworth filter with a bandpass of 0.1-60 Hz. To remove residual BCAs as well as ocular and movement artifacts, all channels were entered into an ICA and decomposed into independent components using the Infomax algorithm. The ICA was run on a copy of the data which was high-pass filtered at 1 Hz to reduce distortion[11]. BCA components were identified based on their topography and Pearson correlation to the EKG segments centered around R peaks in the cardiac cycle. Components reliably categorized as BCAs at the end of this validation procedure as well as components reflecting eye or muscle movement were rejected before back-projecting the ICA to the continuous EEG signal. After data were both GA- and BCA-corrected, they were re-referenced to the average of all scalp electrodes, excluding sensor net positions located on the cheeks or the neck. The continuous data were segmented into five second windows around the presentation of positive and negative feedback (±2500 ms). This was done separately for both T-maze vs. No-maze trials as well as feedback presented in left and right alleys. After all preprocessing was done, three out of the 28 original subjects were excluded due to a lack of valid segments. For the remaining 25 subjects, the standard ERP analysis (see ERP methods above) was performed on the geodesic equivalents of Fz (E8).

MRI data were formatted according to the international Brain Imaging Data Structure[12]. Apart from this, the preprocessing of the functional data was performed identically as in the previous study[13] using custom scripts in MATLAB (release 2020b, Mathworks, Massachusetts, USA) and SPM12. Functional data were first slice-timing and then motion corrected with respect to the first image. Afterwards, functional images were co-registered with the structural scan and then normalized to a standard Montreal Neurological Institute (MNI) template with 12-parameter affine registration. All normalized images were smoothed with an 8-mm$^3$ FWHM Gaussian kernel. First-level analyses were performed using a general linear model (GLM) including a constant, six motion regressors obtained from motion correction, and eight event regressors to model activation time-locked to feedback presentation as well as orientations for the T-maze and No-maze conditions. Events were divided by feedback valence and alley. These four events were further divided by separating by T-maze and No-maze conditions, resulting in four regressors for each alley and feedback[13]. All models were run with a canonical hemodynamic response function and its temporal derivative. Before the preprocessed time series data were statistically modeled, they were high-pass filtered with a mean cutoff period of 128s. To identify voxel activation related to reward effects in the virtual T-maze, all reward feedback was contrasted to no reward feedback.

Finally, linear contrasts of coefficients for each participant were used to run a second level analysis by applying paired t-tests.

**Cortical thickness results:**

In regards to changes in cortical thickness in OU patients compared to healthy controls, four significant clusters of cortical thinning in OU were identified ($p < 0.05$, FDR-corrected): **(1)** left ACC/MCC: nverts = 311, peak vertex xyz = -8, 37, 18 – BA32/24 **(2)** left insula: nverts = 214, peak vertex xyz = -38, 9, -12, **(3)** left posterior cingulate / precuneus region: nverts = 130, peak vertex xyz = -6, -44, 39, and **(4)** left retrosplenial / posterior cingulate cortex: nverts = 159, peak vertex xyz = -3, -46, 16. A main effect of Time or an interaction between Time and Group were not observed.

**EEG-fMRI Results**:

A whole brain contrast between reward vs no reward feedback in the virtual T-maze task yielded five significant reward clusters ($p < 0.05$, FWE-corrected) **(1)** right Putamen: nvoxs = 251 $p < 0.001$, peak voxel: T = 5.85, xyz = 14, -4, -12; **(2)** left Putamen: nvoxs = 732, $p < 0.0001$, peak voxel T = 5.62, xyz = -12, 6, -8; **(3)** left ACC/MCC: nvoxs = 375, $p < 0.0001$, peak voxel: T = 5.09, xyz = -4, 46, 12 - BA32/24; **(4)** left posterior cingulate: nvox = 691, $p < 0.0001$, peak voxel: T = 5.37, voxel xyz = -4, -36, 36 – BA31; **(5)** left Dorsal lateral Prefrontal Cortex: nvoxs = 126, $p < 0.05$, peak voxel: T = 5.1, xyz = -44, 36, 16 – BA46; **(6)** left MCC: nvoxs = 87, peak voxel: T = 4.90, xyz = -2, -12, 34 - BA24.
